# Dissecting the localization of *Tilapia tilapinevirus* in the brain of the experimentally infected Nile tilapia (*Oreochromis niloticus*)

**DOI:** 10.1101/2021.01.19.427199

**Authors:** Nguyen Dinh-Hung, Pattiya Sangpo, Thanapong Kruangkum, Pattanapon Kayansamruaj, Tilladit Rung-ruangkijkrai, Saengchan Senapin, Channarong Rodkhum, Ha Thanh Dong

## Abstract

*Tilapia tilapinevirus* or tilapia lake virus (TiLV) is an emerging virus that inflicts significant mortality on farmed tilapia globally. Previous studies reported detection of the virus in multiple organs of the infected fish; however, little is known about the in-depth localization of the virus in the central nervous system. Herein, we determined the distribution of TiLV in the entire brain of experimentally infected Nile tilapia. *In situ* hybridization (ISH) using TiLV-specific probes revealed that the virus was broadly distributed throughout the brain. The strongest positive signals were dominantly detected in the forebrain (responsible for learning, appetitive behavior, and attention) and the hindbrain (involved in controlling locomotion and basal physiology). The permissive cell zones for viral infection were observed mostly to be along the blood vessels and the ventricles. This indicates that the virus may productively enter into the brain through the circulatory system and widen broad regions, possibly through the cerebrospinal fluid along the ventricles, and subsequently induce the brain dysfunction. Understanding the pattern of viral localization in the brain may help elucidate the neurological disorders of the diseased fish. This study revealed the distribution of TiLV in the whole infected brain, providing new insights into fish-virus interactions and neuropathogenesis.

## 1. Introduction

An unknown viral disease associated with abnormal mortalities (>80%) of farm-bred tilapia Chitralada during 2011-2012 was first reported in Ecuador in May 2013. The disease was initially named syncytial hepatitis of tilapia (SHT) based on the pathognomonic lesion in the liver (Ferguson et al., 2014). At the same time, similar mysterious disease episodes occurred in Israel and a novel virus was subsequently discovered and initially termed tilapia lake virus (TiLV) (Eyngor et al., 2014). The causative viruses from these cases were later confirmed to be identical (Bacharach et al., 2016; Del-Pozo et al., 2017). Since then, the virus has been recognized as an emerging virus causing disease problems in 16 countries across the continents of Asia, Africa, and South America and severely impacts the tilapia industry (FAO, 2018; Jansen, Dong, & Mohan, 2019; OIE, 2017). The virus was initially described as a novel *Orthomyxo-like* virus with a 10 negative sense, single-stranded RNA segmented genome (Bacharach et al., 2016; Eyngor et al., 2014) and was later classified as *Tilapia tilapinevirus*, a single species belonging to the *Tilapinevirus* genus, under the *Amnoonviridae* family (ICTV, 2019). However, TiLV remains a common name in both scientific and non-scientific documents.

Tilapia lake virus is a severe contagious pathogen that causes high mortality (20-90%) in both farmed and wild tilapines (Al-Hussinee et al., 2019; Bacharach et al., 2016; Dong et al., 2017; Eyngor et al., 2014; Ferguson et al., 2014; Jansen et al., 2019). The infected fish exhibited variable gross signs i.e. skin erosion, scale protrusion, or gill pallor to abdominal distension, anemia, exophthalmia as well as abnormal behaviors. The latter included lethargy, loss of appetite, swimming at the water surface, erratic swimming, stoppage of schooling, swirling or loss of balance (Bacharach et al., 2016; Dong et al., 2017; Fathi et al., 2017; Ferguson et al., 2014; Jansen et al., 2019; Surachetpong et al., 2017; Tattiyapong, Dachavichitlead, & Surachetpong, 2017). These unusual behaviors may have involved damages within the central nervous system. The previous studies have demonstrated various histopathological changes in the brain of infected fish notably as congestion of blood vessels and perivascular cuffing of lymphocytes (Behera et al., 2018; Eyngor et al., 2014; Fathi et al., 2017; Ferguson et al., 2014; Jansen et al., 2019). Furthermore, the presence of the TiLV genomic RNA in the infected brain has also been determined by using *in situ* hybridization (Bacharach et al., 2016; Dong et al., 2017). However, these findings were limited at suggesting that the brain is one of the target tissues for transcription and replication of the virus. In fact, the fish brain divides into different regions and each part is responsible for certain functions (Table 1) (Baldisserotto, Urbinati, & Cyrino, 2019; Roberts, 2012). Unfortunately, currently available data is unclear as to which part of the brain is affected and whether the possible injuries are linked to the brain function, as well as the possible route of viral entry into the brain. Therefore, elucidating the localization of the virus in the brain is an important background for better understanding the neuropathogenesis induced by TiLV infection. Thus, the present study investigated the spatial localization of TiLV in the entire infected brain detected by ISH using the two different specific probes. Additionally, brain histopathological changes caused by TiLV are also described. In drawing things to a close, the potential route of TiLV to the fish brain was suggested. To our best knowledge, this is the first in-depth study of the distribution of TiLV and its possible effects in infected fish brains, in cooperation with the histopathological study, providing an initial step for further understanding the disease manifestation and host-pathogen interaction.

**TABLE 1.** Distribution of TiLV signals in the brain of infected fish and possible link to abnormal behaviors

| Brain region |  | Abbreviation | Densities of signal | Brain Function* | Possible link to behavioral abnormalities |
| --- | --- | --- | --- | --- | --- |
| Forebrain | Telencephalon |  |  | Learning, appetitive behavior and attention | Loss of appetite, lethargy and stoppage of schooling |
|  | Olfactory bulb | OB | +++ |  |  |
|  | Hemisphere of telencephalon | HT | +++ |  |  |
|  | Periventricular zone of telencephalon | PZT | +++ |  |  |
|  | Diencephalon |  |  | Homeostatic and appetitive coordination | Losing appetite and stopping eating |
|  | Thalamus | Tha | + |  |  |
|  | Optic nerve | ON | + |  |  |
|  | Hypothalamus | Hyp | ++ |  |  |
|  | Diffuse nucleus of inferior lobe | DIL | + |  |  |
|  | Periventricular zone of hypothalamus | PZH | +++ |  |  |
| Midbrain | Mesencephalon |  |  | Reception and coordination of optic nerve inputs | Disorders of navigating, food and prey seeking |
|  | Optic tectum | TeO | + |  |  |
|  | Periventricular gray zone of optic tectum | PGZ | ++ |  |  |
|  | Tegmentum | Teg | + |  |  |
|  | Torus longitudinalis | TL | ++ |  |  |
|  | Torus semicircularis | TS | + |  |  |
| Hindbrain | Metencephalon |  |  | Regulation of locomotion and balance stimuli | Erratic swimming or loss of balance |
|  | Corpus division of cerebellum | CCe | ++ |  |  |
|  | Granular zone of corpus cerebellum | CCeG | ++ |  |  |
|  | Molecular zone of corpus cerebellum | CCeM | + |  |  |
|  | Valvula division of cerebellum | VCe | + |  |  |
|  | Medulla oblongata |  |  | Basal physiology (regulation of the respiratory and cardiovascular systems) | Result in high mortality |
|  | Intermediate reticular formation | IMRF | + |  |  |
|  | Inferior reticular formation | IRF | + |  |  |
|  | Vagal lobe (lobe of nerve X) | LX | +++ |  |  |

(ISH signals: + + +, *high*; + +, *moderate*; +, *low*; −, *absent*)
\*according to Roberts, 2012; Baldisserotto et al., 2019

## 2. Materials and methods

### 2.1. Viral preparation

Tilapia lake virus strain NV18R isolated from diseased hybrid red tilapia (*Oreochromis* sp.) using E11 cell line (Dong et al., 2020) was used for challenge test in this study. Prior to the experiment, the viral stock (10^7.5^ TCID_50_ per mL) preserved at −80 °C was thawed and diluted 1:10 with saline solution (0.9% NaCl) to be used as an inoculum dose of 10^5.5^ TCID_50_ per fish.

### 2.2. Experimental setup

Apparently normal Nile tilapia (*Oreochromis niloticus*) fingerlings (10 g ± 1 g body weight) were obtained from a tilapia hatchery with no history of TiLV infection. The fish were acclimatized in a laboratory rearing facility for 10 days and water temperature was set at 28°C ± 1°C. Prior to challenge test, ten fish were randomly selected for the detection of TiLV by semi-nested RT-PCR (Dong et al., 2017) to warrant TiLV negative status. These fish were divided into three groups with 20 individuals in a 50 L fiberglass tank. Each tank was provided with a separate water supply, drainage, and air stones. The tilapia fish were fed with commercial tilapia feed pellets (Charoen Pokphand, Thailand) containing 28% crude protein twice daily (9.00 am and 16.00 pm) at 3% biomass per day. Every three days, 50% of the water in each tank was replaced. The fish in group 1 and group 2 were intraperitoneally injected with 0.1 mL TiLV to receive a dose of 10^5.5^ TCID_50_ per fish, while the fish in the group 3 were injected with 0.1 mL of normal saline solution as control.

### 2.3. Clinical observation and brain collection

Following experimental infection, the fish in group 1 were observed continuously for 16 days to record gross pathology and mortality. While, in the group 2, moribund fish that showed clear signs, which prioritized neurological manifestation such as loss of appetite, lethargy and abnormal behavior (e.g. swimming at the surface, stoppage of schooling, erratic swimming, or loss of balance), were euthanized with an overdose of (150 ppm) of clove oil before necropsy. The intact brain of diseased fish (n = 6) were carefully isolated and preserved in 10% neutral buffered formalin for 24 hr and then immersed in 70% ethanol before being processed for routine histology and *in situ* hybridization, as described below.

### 2.4. Tissue processing and histopathology

The preserved tissues were dehydrated by incubating in several increasing concentrations of ethanol (70–100%) and then transferred to xylene automatically by Leica TP1020 Tissue Processor (Leica Biosystems, US). The tissues were infiltrated and embedded in paraffin. Each paraffin-embedded tissue was sequentially sectioned at 5 μm thickness into 5 consecutive sections and were mounted on HistoGrip (Invitrogen, US) coated glass slides. The purposed tissue sections included 2 slides for ISH with TiLV specific probes derived from TiLV genome segment 1 and segment 3 (see below), 2 slides for ISH with negative controls (unrelated probe and no probe), and one slide for hematoxylin and eosin (H&E) staining. To analyze the distribution of TiLV in the whole-brain, the samples were sectioned toward horizontal and parasagittal sides and the results were interpreted in parallel.

### 2.5. *In situ* hybridization (ISH) assay

#### 2.5.1. Preparation of probes

Two TiLV-specific DIG-labelling probes targeting two different TiLV genomic segments were used for comparison and double confirmation of viral localization in this study. The 274 bp probe derived from TiLV segment 1 was prepared using primers TiLV/nSeg1F; 5’-TCT GAT CTA TAG TGT CTG GGC C-3’ and TiLV/nSeg1RN; 5’-CCA CTT GTG ACT CTG AAA CAG–3’ (Taengphu et al., 2020) while the 250 bp probe derived from TiLV segment 3 employed primers ME1; 5’-GTT GGG CAC AAG GCA TCC TA-3’ and 7450/150R; 5’-TAT CAC GTG CGT ACT CGT TCA GT-3’ (Eyngor et al., 2014; Tsofack et al., 2017). An unrelated probe 282 bp acquired from a shrimp virus namely infectious myonecrosis virus (IMNV) (F13N; 5’-TGT TTA TGC TTG GGA TGG AA-3’ and R13N; 5’-TCG AAA GTT GTT GGC TGA TG-3’) (Senapin, Phewsaiya, Briggs, & Flegel, 2007) and no probe were used as negative controls. The probes were prepared using Digoxigenin (DIG), a commercial PCR DIG-labeling mix (Roche Molecular Biochemicals, Germany). Briefly, RNA extracted from internal organs of tilapia infected with TiLV were used as a template for one step of RT-PCR to amplify 274 bp and 250 bp of genomic segments 1 and 3 as described previously (Dong et al., 2017; Taengphu et al., 2020). The targeted fragments were cloned into pGEM^®^-T Easy (Promega, US). The plasmids containing TiLV segments 1 and 3, were then used as templates for labeling reaction with DIG by PCR. The PCR reaction of 25 μL was composed of 200 ng of plasmid template, 1 μM of each primer, 0.5 μL of Platinum^®^ Taq polymerase (Invitrogen, US), and 1× of the supplied buffer, dNTP in the reaction was replaced by 0.5 μL DIG-labeling mix. The reactions were heat activated at 94°C for 5 min. PCR cycling then was carried out for 30 cycles at 94°C for 30 s, 55°C for 30 s, 72 °C for 30 s, final extension step at 72 °C for 5 min. The DIG-labeled TiLV probes for each segment was obtained after purifying amplified product by NucleoSpin™ Gel PCR Clean-up Kit (Fisher Scientific, US) according to the manufacturer’s protocol. The purified probes were measured for concentration by NanoDrop™ One Spectrophotometer (Fisher Scientific, US) and stored at −20 °C until use.

#### 2.5.2. Prehybridization and hybridization

Unstained 5 μm sections on HistoGrip coated slide was deparaffinized three times in xylene for 5 min followed by a graded series of ethanol (95%, 80%, 75% twice each for 5 min), distilled water, and finally in TNE buffer (100 mM Tris–HCl, 10 mM EDTA, pH 8.1). Tissues were digested with proteinase K (prepared just prior to use) at a final concentration of 10 μg mL^−1^ for 15 min at 37°C, then treated by 4% paraformaldehyde for 5 min at 4°C and immersed in distilled water for 5 min. After rapidly treating with acid acetic for 20 s and washing in distilled water, each section was then covered with pre-hybridization buffer (4× SSC containing 50% (v/v) deionized formamide) at 37 °C for at least 10 min. Each probe was diluted in hybridization buffer (50% deionized formamide, 50% dextran sulfate, 50× Denhardt’s solution (Sigma, Germany), 20 × SSC, 10 mg mL^−1^ salmon sperm DNA (Invitrogen, US), heated at 95°C for 10 min, and then chilled on ice. Each specific probe was added to the tissue sections, then covered by coverslips, and incubated overnight at 42°C in a humid chamber.

#### 2.5.3. Post-hybridization

Post-hybridization was performed by sequentially washing twice each, with 2×, 1× and 0.5× SSC at 42°C, 37°C, 37°C for 15 min, respectively, equilibrated by 5 min washing with buffer I (1 M Tris-HCl, 1.5 M NaCl, pH 7.5). Tissue sections were then blocked with blocking solution buffer II (containing 0.1% Triton X-100 and 2% normal sheep serum) at room temperature for 30 min before covering with anti-DIG alkaline phosphatase conjugate anti-digoxigenin antibody (Roche, diluted 1:500 in buffer II) for 1 h at 45°C. After washing twice for 10 min each with Buffer I, the sections were treated for 10 min in buffer III (100 mM Tris–HCl, 1.5 M NaCl, 50 mM MgCl2, pH 9.5). The sections were incubated for 1 to 24 hr in the dark at room temperature with development solution (NBT/BCIP substrate, Roche)

#### 2.5.4. Detection and visualization

The antibody-antigen complexes were subsequently revealed by NBT/BCIP substrate. Once the optimal color was observed the reaction was then stopped by washing tissue sections with 1× TE buffer at room temperature for 15 min and afterward dripped in distilled water. The slides were then counterstained with 0.5% Bismarck Brown (Sigma Aldrich, US) for 2 min. The sections were washed under running water for 5 min then dried at room temperature before immersing twice in 100% xylene for 5 min each. The slides were then mounted, coverslipped, observed and photographed under a light microscope BX51 (Olympus, Japan). Localization of TiLV was interpreted in parallel with the slides of TiLV-specific probes, negative controls (unrelated probe and no probe), and H&E stained sections. TiLV signal density in the brain was subjectively scored on a four-point scale as follows: + + + (*high signal*), + + (*moderate signal*), + (*low signal*), and – (*absent signal*) following Cham et al (2017). Nomenclatures for the brain area (Table 1) were based on those described by Wullimann et al (1996), Simoes et al (2012), Ogawa et al (2016), and Cham et al (2017).

## 3. Results

### 3.1. Clinical observations and cumulative mortality

After experimental infection, the fish showed no remarkable abnormalities in appetitive behavior as well as swimming activity during the first two days. Starting at 3 days post-infection (dpi), some of the fish showed lost of appetite, separated from the group, and swimming near the water surface. At 4 dpi, some fish became lethargic and stopped eating. Mortality started at 5 dpi and lasted until 11 dpi with cumulative mortality at 95% (Figure 1A). Prior to death, approximately 20% of fish displayed erratic swimming, swirling, or loss of balance during 7-10 dpi (Figure S1). A majority of the fish died within 24 hr after appearance of abdominal swelling, exophthalmia and dark discoloration of the skin (Figure 1B). In addition, some sick fish also displayed scale protrusion and skin erosion (data not shown). Internally, post-mortem changes including necrotic and pale liver; and enlarged spleen were frequently noticed as well as ascitic fluid was observed (Figure 1C). In contrast, no clinical signs of infection or mortality were observed in the control group.

**FIGURE 1.**
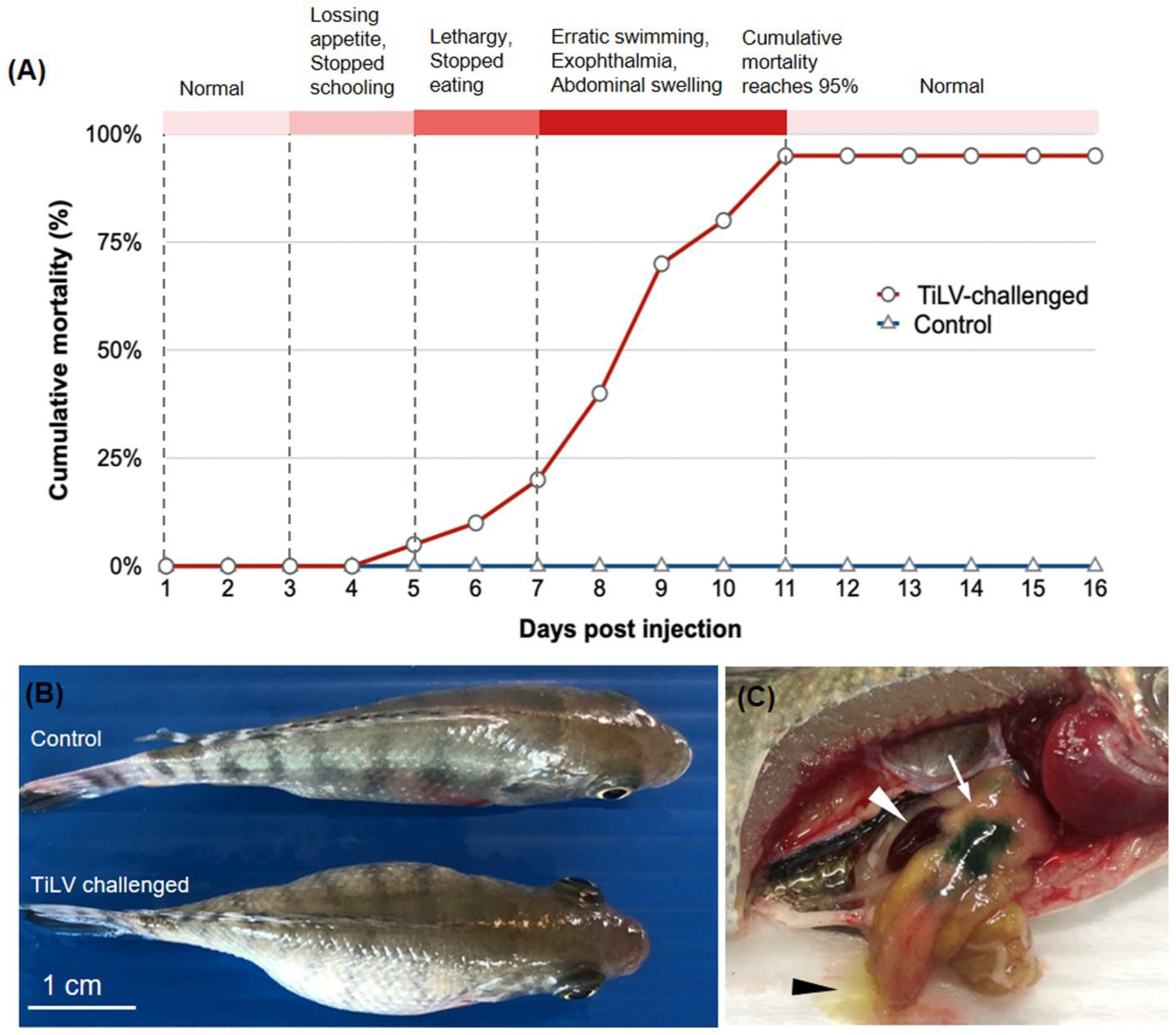
Experimental challenge of Nile tilapia with TiLV. (A) Clinical signs and cumulative mortality of Nile tilapia following injection (i.p.) with TiLV at 10^5.5^ TCID_50_/fish and the control were injected (i.p.) with 0.1 mL of 0.9% saline solution. (B) Diseased fish showed clinical signs of exophthalmia and abdominal swelling. (C) Necrotic and pale liver diffused with green bile (white arrows), enlarged spleen (white arrowhead) as well as ascitic fluid (black arrowhead).

### 3.2. TiLV Localization in the fish brain

The results of ISH using two specific probes targeting TiLV genome segment 1 and segment 3 exhibited similar localization of TiLV positive signals but difference in intensities. In contrast, no signal was detected in the sections from the same samples assayed with an unrelated probe and no probe. ISH positive signals (dark color) were widely distributed in various parts of the brain. However, the forebrain and hindbrain showed higher signal densities and stronger signal intensities compared to that of the midbrain. Details on the distribution of the TiLV positive signals and their densities are described below and summarized in Table 1. Representative microphotographs of the horizontal and parasagittal whole-brain from diseased fish are shown in Figure 2 and Figure 3, respectively. Results were read from four consecutive sections of the fish brains subjected to H&E staining and ISH with either segment 1 probe, segment 3 probe or unrelated probe. There were no ISH positive signals detected in the brain of non-infected control fish that were assayed in the same manner (Figure S2).

**FIGURE 2.**
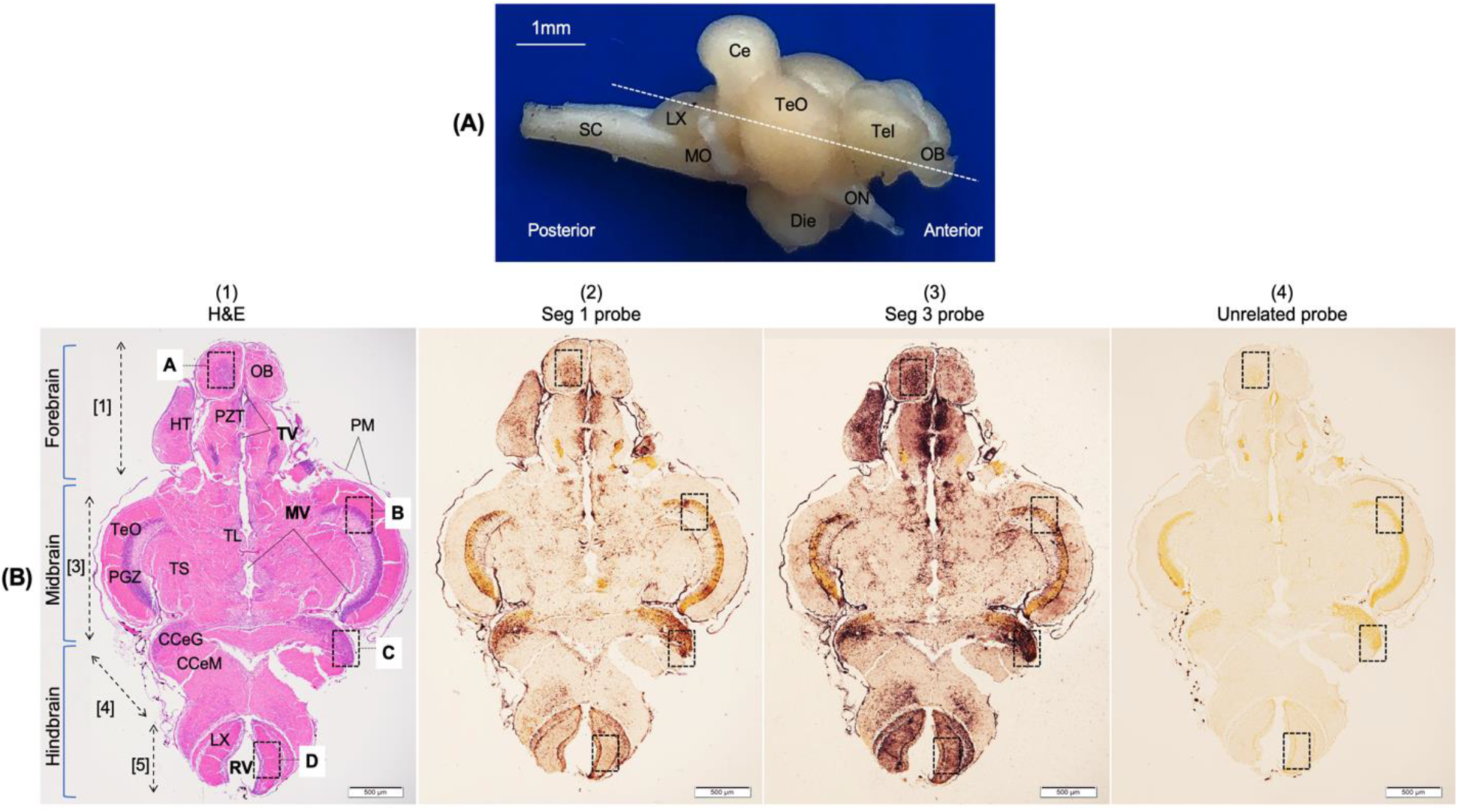
The spatial localization of TiLV in the brain of infected fish (horizontal sections). (A) Lateral view of tilapia brain with approximate slice position (white dashed line). (B) Comparison of horizontal consecutive sections from infected fish stained with H&E (B.1), ISH with TiLV probe prepared from genomic segment 1 (B.2), genomic segment 3 (B.3) and ISH with unrelated probe (B.4) as the control. Positive reactivity is shown by dark brown signals. The horizontal section was divided four major areas: telencephalon [1], mesencephalon [3], metencephalon [4], medulla oblongata [5]. Dotted boxes (A, B, C, D) are exhibited at the high magnification images in Figure 6. Ce: Cerebellum, Die: Diencephalon, MO: Medulla oblongata, MV: Mesencephalic ventricle, Tel: Telencephalon, TV: Telencaphalic ventricle, RV: Rhombencephalic ventricle, SC: Spinal cord. Other abbreviations are listed in Table 1. Specimen was collected on 8 dpi. The scale bars are shown in the pictures.

**FIGURE 3.**
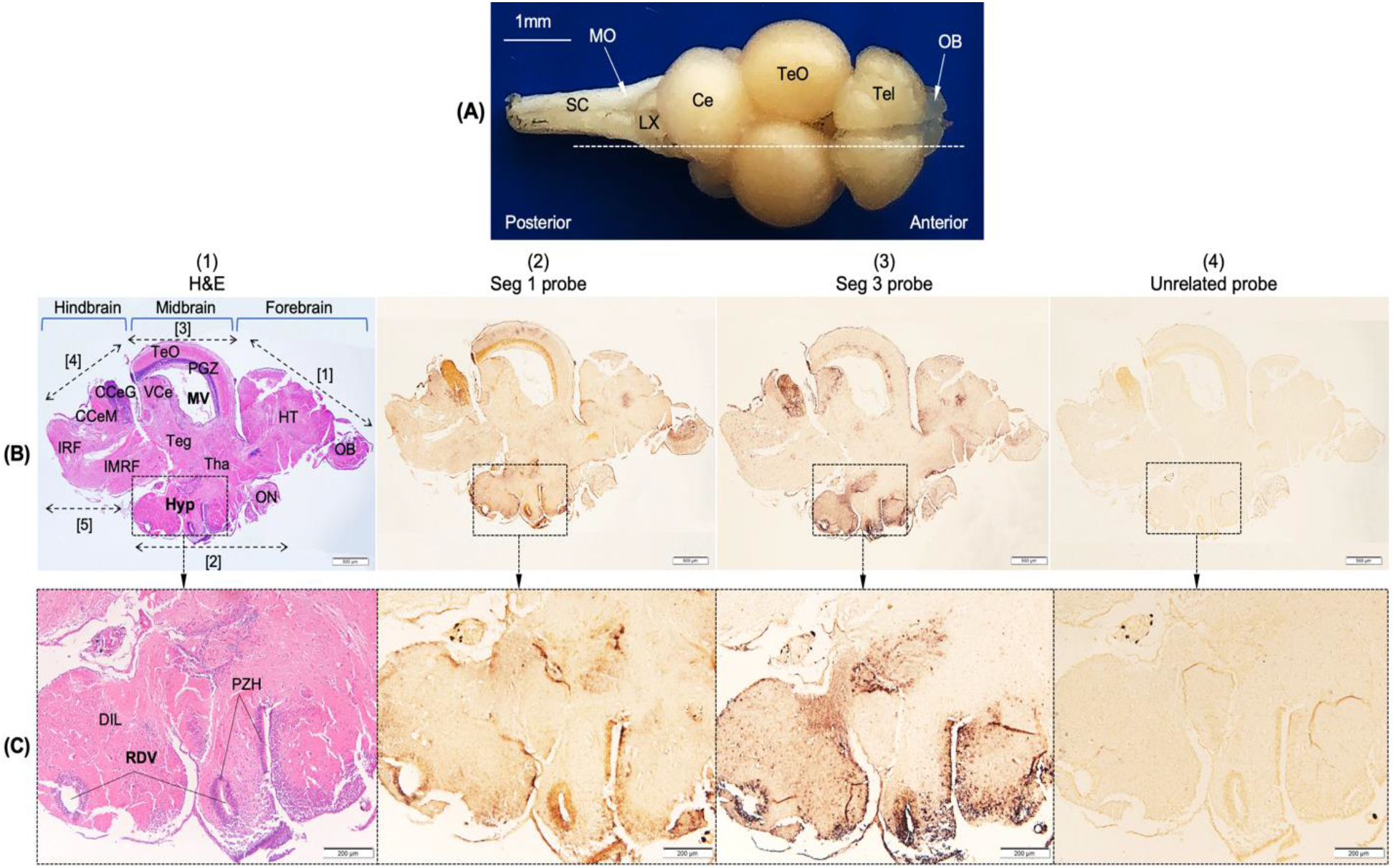
The spatial localization of TiLV in the brain of infected fish (parasagittal sections). (A) Dorsal view of tilapia brain with approximate slice position (white dashed line). (B) Comparison of parasagittal consecutive sections from infected fish stained with H&E (B1), ISH with TiLV probe based on genome segment 1(B2), genome segment 3 (B3), and ISH with unrelated probe (B4) as the control. (C) Higher magnification of dotted boxes in (B). The parasagittal section allows visualization of five major areas: telencephalon [1], diencephalon [2], mesencephalon [3], mesencephalon [4], medulla oblongata [5]. Ce: Cerebellum, MO: Medulla oblongata, MV: Mesencephalic ventricle, Tel: Telencephalon, RDV: Recess of diencephalic ventricle, SC: Spinal cord. Other abbreviations are listed in Table 1. Specimen was collected on 9 dpi. The scale bars are shown in the pictures.

#### The forebrain (prosencephalon)

The forebrain comprises of two main parts, namely telencephalon (or cerebrum) and diencephalon (approximately defined by areas [1] and [2] in Figure 2B.1 and 3B.1). In the telencephalon [1], all subdivisions; the olfactory bulbs (OB), hemispheres of telencephalon (HT), and periventricular zone (PZT), were positively reacted with the TiLV probes. OB and HT had more intense signals in the granular cell layer while dense positive staining was observed in a large number of cells localized in the PZT (Table 1, Figure 2B.2, 2B.3, 3B.2 and 3B.3). In the diencephalon [2], there were relatively few positively labelled cells in the thalamus, optic nerve, and diffuse nucleus of the inferior lobe. Strong positive staining was found throughout the hypothalamus (Hyp), specifically along the periventricular zone of hypothlamus (PZH) (Table 1, Figure 3C.2 and 3C.3). Note that signals from the probe prepared from TiLV genomic segment 3 were stronger than that from segment 1 probe.

#### The midbrain (mesencephalon)

The midbrain or mesencephalon (marked as area [3] in Figure 2B.1 and 3B.1) divides mainly into parts including optic tectum (TeO), torus longitudinalis (TL), torus semicircularis (TS), tegmentum (Teg), and periventricular grey zone of the optic tectum (PGZ) (Figure 2B.1 and 3B.1). Diffuse staining of positively TiLV labelled cells was observed in TeO and TS. Stronger signals were observed in PGZ and TL (Table 1, Figure 2B.2, 2B.3, 3B.2 and 3B.3). In the Teg, positive signals were weaker compared to other areas (Figure 3B.2 and 3B.3).

#### The hind brain (rhombencephalon)

The hindbrain consists of metencephalon (or cerebellum) and medulla oblongata (areas [4] and [5], respectively in Figure 2B.1 and 3B.1). There were relative stronger positively TiLV labelled cells in the hindbrain compared to the midbrain. Within the cerebellum [4], positive staining signals were abundantly observed in the granular layers of the corpus cerebellum (CCeG) (Table 1, Figure 2B.2, 2B.3, 3B.2 and 3B.3). In the medulla oblongata [5], TiLV signals were densely localized in the vagal lobe (LX) (Figure 2B.2 and 2B.3). Fewer TiLV-positive signals were seen in the molecular zone of corpus cerebellum (CCeM), intermediate reticular formation (IMRF), and inferior reticular formation (IRF) (Figure 2B.2, 2B.3, 3B.2 and 3B.3).

### 3.3. The permissive cell zones and pattern of TiLV distribution area

Although the viral positive signals were presented throughout the brain, ISH revealed strong positive signals of TiLV infected location in the primitive meninges (PM) (Figure 4) and in the periventricular regions (Figure 5) (both areas close to cerebrospinal fluid). Additionally, the heavy ISH signals were also detected in the epithelium of the blood vessels (arrowheads in Figure 4A, 4B, and 5C). In the periventricular region, the signal intensity indicated a distinctive pattern with a gradual decrease from the strongest signals localizing close to the ventricle to lesser infected cells in the area farther away from the ventricle (see Table 1). In particular, TiLV signals were detected in the ependymal cells lining the ventricles and in the choroid plexus epithelial cells (Figure 5).

**FIGURE 4.**
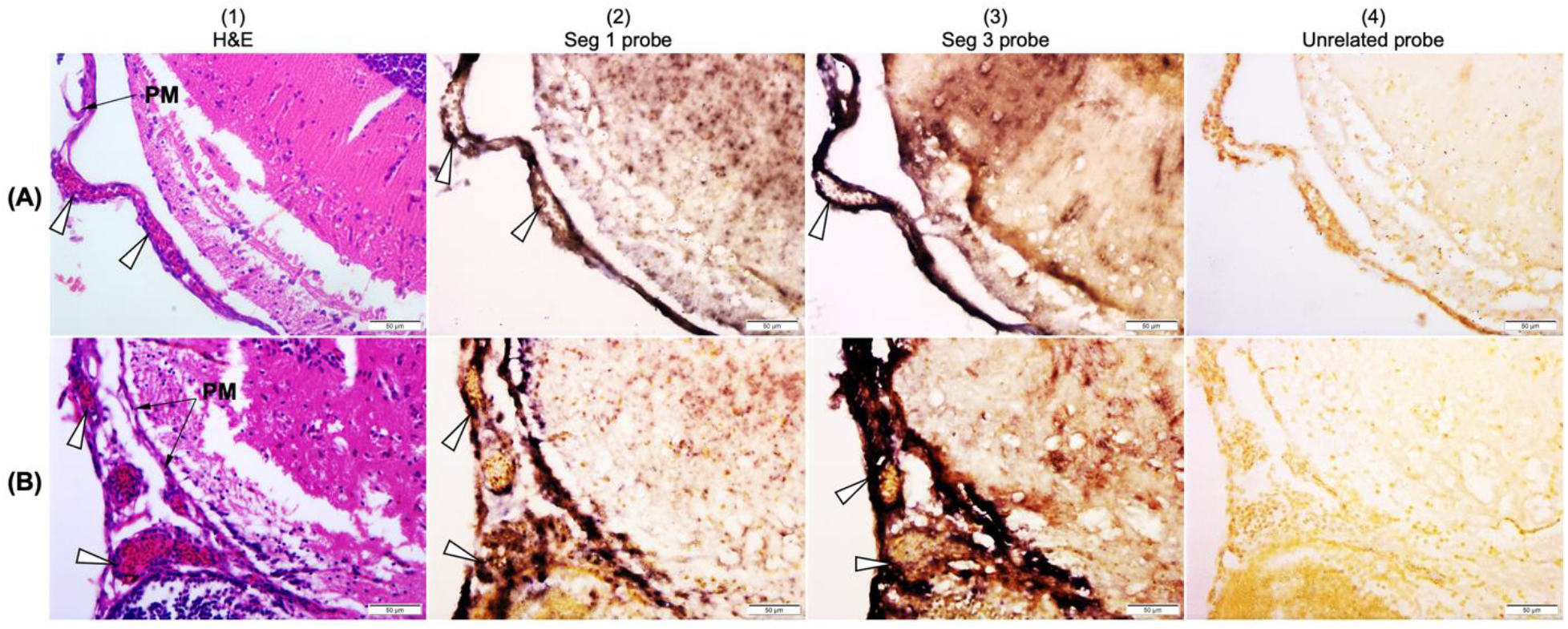
Microphotographs revealed localizations of TiLV in the brain blood vessels and primitive meninges (PM). H&E staining sections, ISH with TiLV-specific probe, and ISH sections with unrelated probe of two representative TiLV infected fish brains are shown in (A) and (B). Viral RNA (dark staining) was strongly detected in the primitive meninges (PM) and the endothelial cells of the congested blood vessels (arrowhead). Specimen was collected on 8–9 dpi. The scale bars are shown in the pictures.

**FIGURE 5.**
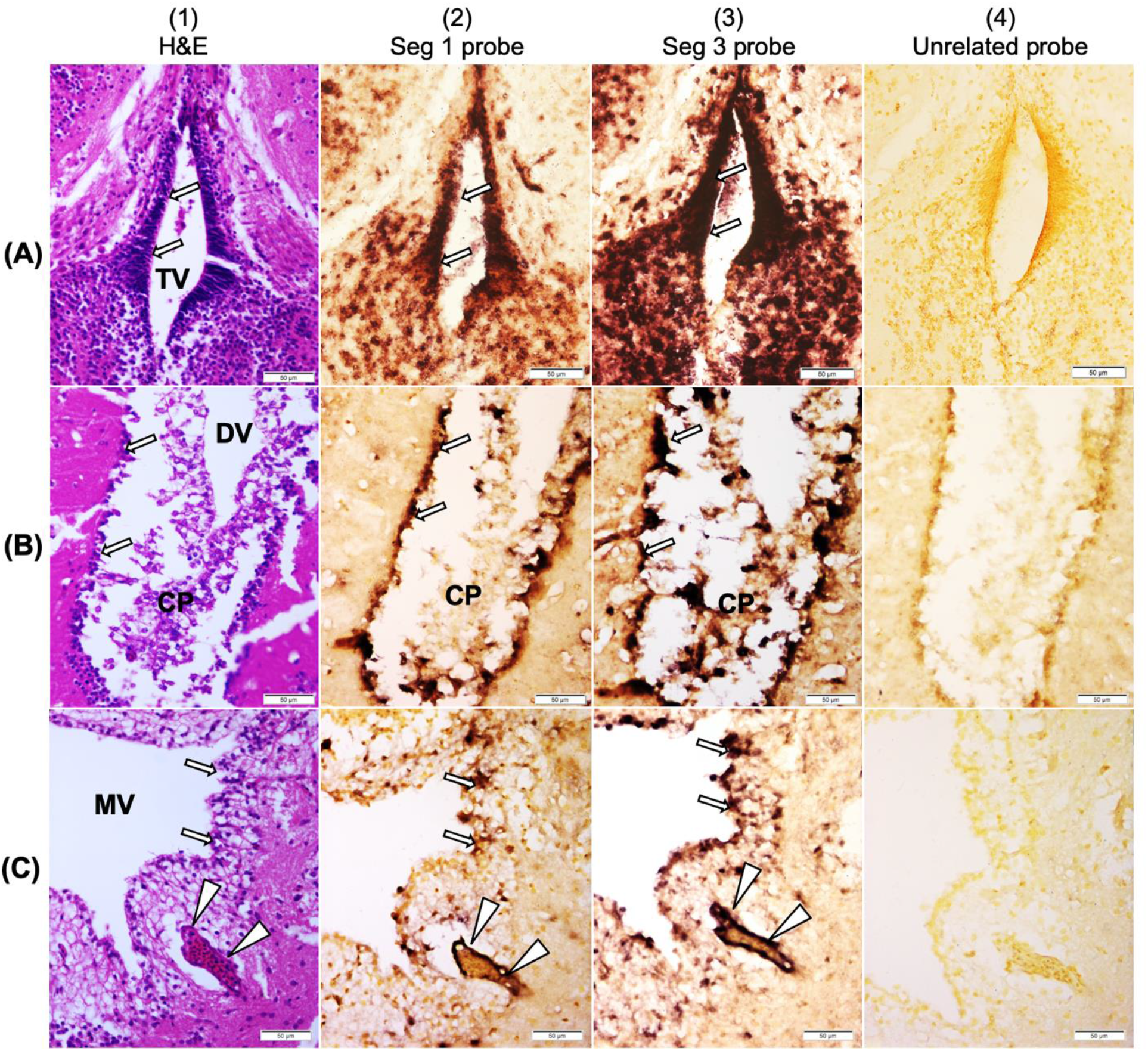
*In situ* hybridization for detection of TiLV in the brain (ventricles and choroid plexus). H&E staining sections, ISH with TiLV-specific probe, and ISH sections with unrelated probe of three representative TiLV infected brains are shown in (A-C). Viral RNA (dark staining) was detected in the periventricular regions including the ventricular ependymal cells (arrows), choroid plexus (CP) and endothelial cells of the blood vessels (arrowheads). DV: Diencephalic ventricle, MV: Mesencephalic ventricle, TV: Telencaphalic ventricle. Specimen was collected on 8-9 dpi. The scale bars are shown in the pictures.

### 3.4. Histopathological assessment of the infected brain

Examination of histopathological alterations in correlation with TiLV positive signals in the infected brain revealed the presence of cells dissociation and degeneration within the infected sites including the internal cellular layer of the olfactory bulb in the telencephalon (Figure 6A), periventricular grey zone (PGZ) of the optic tectum in the mesencephalon (Figure 6B), granular layer of the corpus cerebellum (Figure 6C), the motor layer, sensory layer and fiber layer of the vagal lobe in the medulla oblongata (Figure 6D). No apparent histological alterations were visible in the respective areas of the normal brain (first column panel in Figure 6). In addition, we found some other obvious histopathology alterations in various areas of the infected brain such as inflammation of meninges (Figure 4B.1), congestion of blood vessels (Figure 4A.1, 4B.1, 7A and 7B) and cell aggregation (Figure 7C). Cell aggregation can be found in the granular layer of the cerebellum and the olfactory bulb.

**FIGURE 6.**
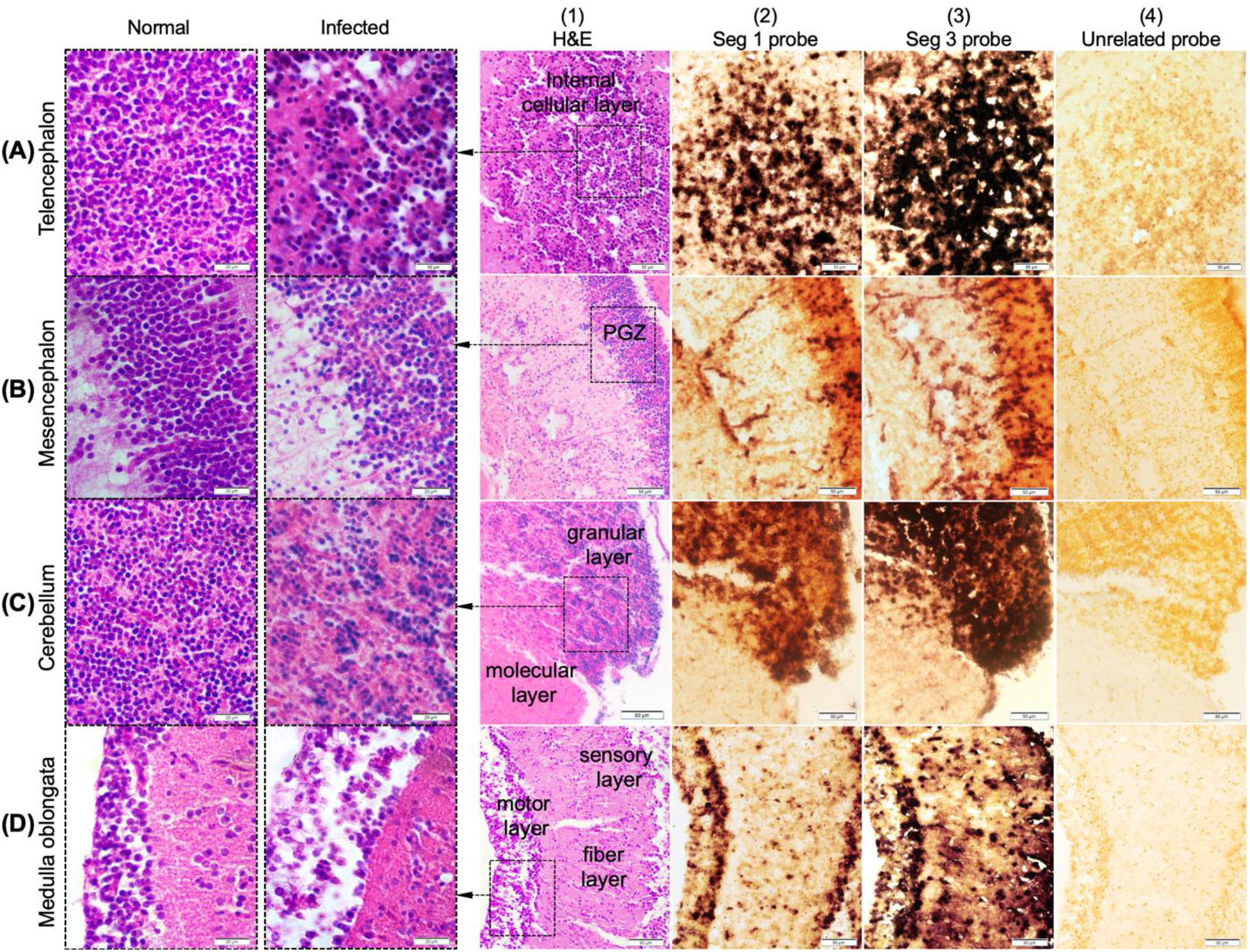
Representative higher magnification photomicrographs from Figure 2 showing TiLV distribution in different parts of the brain of infected tilapia. Strong TiLV positive signals were detected in the internal cellular layer (ICL) of the olfactory bulb in the telencephalon (A); periventricular grey zone (PGZ) of optic tectum in the mesencephalon (B); granular zone of the corpus cerebellum (CCeG) (C); motor layer, sensory layer and fiber layer of the medulla oblongata (D). Cells dissociation and degeneration were visible within the location that showed the TiLV positive signals in H&E stained sections (third column panel and higher magnification in second column panel). H&E stained respective brain locations of normal fish (first column panel) were also included for comparison. The scale bars are shown in the pictures.

**FIGURE 7.**
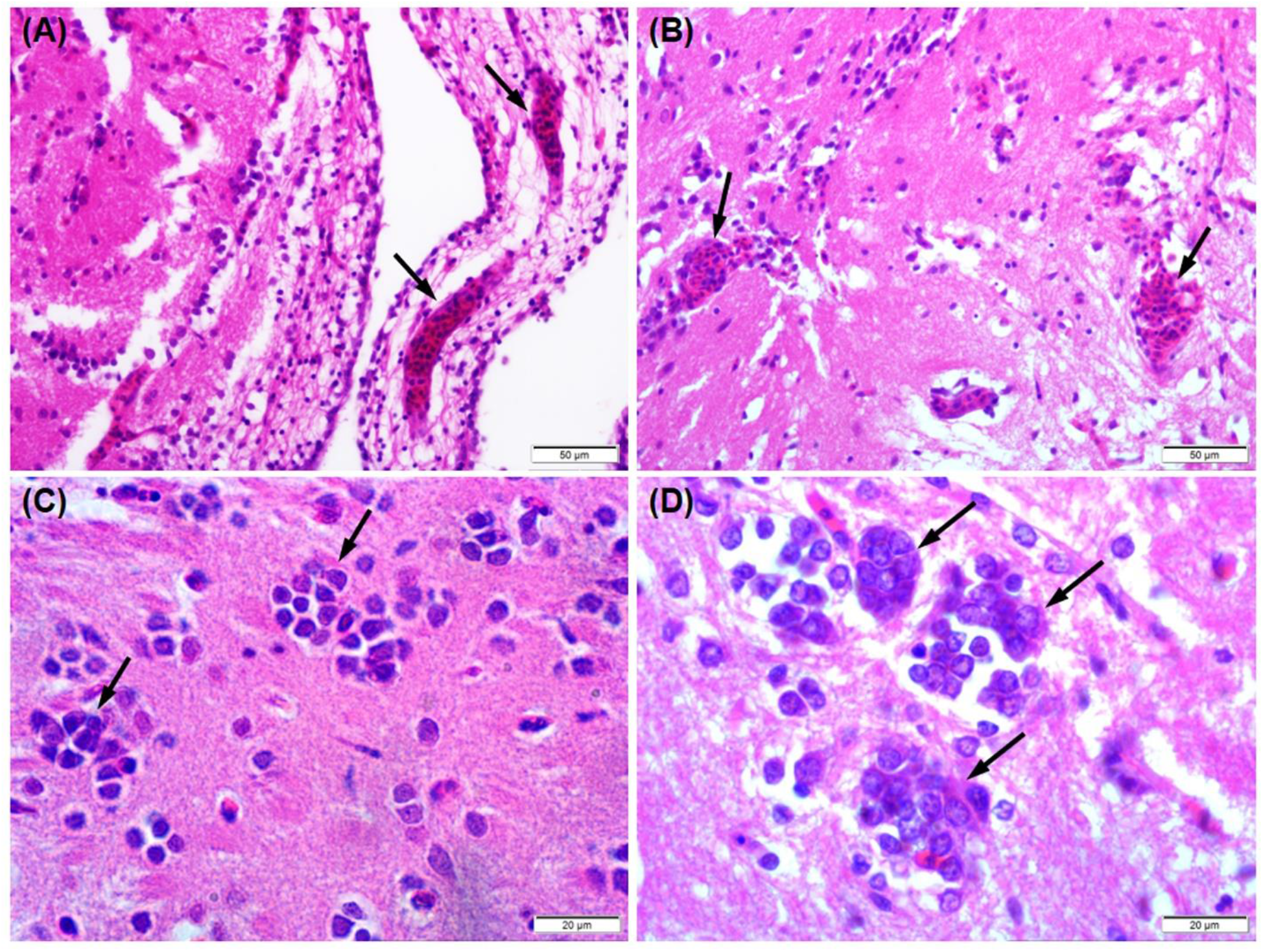
Photomicrographs of histopathological alterations in the TiLV infected brain. Congestion of blood vessels (arrow in A and B) and cell aggregation (arrows in C and D) was found in various areas. The scale bars are shown in the pictures.

## 4. Disccussion

Localization of TiLV was initially investigated by ISH in a previous study, however, only one part of the brain resembling to optic tectum of the midbrain was explored (Dong et al., 2017). In this study, the localization of TiLV in the whole-brain of infected Nile tilapia was comprehensively dissected for the first time. Distribution of positive signals was detected throughout the brain, however, the viral positive signals appeared to be more concentrated in some particular areas of the forebrain and hindbrain. These findings suggest that these regions possibly contain more permissive cells for propagation of TiLV than that of the midbrain. With respect to disease diagnosis, these findings imply that the fore- and hindbrain might be the target tissues with predominant of the virus. It should be noted that, when two TiLV-specific probes targeting different genome segments of TiLV in parallel for ISH assay were used, similar distribution of the positive signals double confirmed the localization of the virus in major parts of the brain. However, minor differences in signal distribution and intensities may possibly be derived from frequency of Digoxigenin-11-dUTP incorporated into newly synthesized DNA probes. It is also possible that relatively higher %GC content and melting temperature (Tm) of segment 3 probe (52.0% GC, Tm 83.5 °C) makes it more effective in hybridization reaction than that of segment 1 probe (48.5% GC, Tm 82.3 °C).

Despite the fact that this study did not focus on functional study of the fish brain, basic science revealed that the brain is a central nervous system that controls important living activities of the fish and different regions of the brain are responsible for different biological functions (Table 1) (Baldisserotto et al., 2019; Ferguson, 2006; Northcutt, 1981; Northcutt, 1995; Roberts, 2012). Therefore, heavy viral infection (indicated by density of ISH signals) in particular brain regions may result in neurological cell damages and impairment of the brain function and thus possibly explain for abnormal behaviors observed in infected fish during the course of infection. The abnormal behaviors such as lethargy, loss of appetite, erratic swimming, stoppage of schooling were consistently recorded in this study and several previous studies (Table 1) (Dong et al., 2017; Tattiyapong et al., 2017). Indeed, based on literature, the teleost brain regions in the fore-, mid-, and hindbrain are involved with specific physiological and behavioral outputs. Specifically, the forebrain contains the telencephalon and the diencephalon in which the telencephalon encompasses control of sensory and motor as well as cognitive tasks like memory, learning, and emotion (Table 1) (Baldisserotto et al., 2019; Northcutt, 1981; Northcutt, 1995; Roberts, 2012). Heavy TiLV infection in this important brain part could thus result in some described symptoms such as lethargy, stoppage of schooling, and swimming at the water surface. The diencephalon function mainly as correlation centers for sensory inputs such as gustation and olfaction (Ferguson, 2006; Muñoz-Cueto & College, 2001). With respect to the symptoms of loss of appetite and stoppage of eating, it is interesting that the intense TiLV signals were observed in the hypothalamus, which controls feeding behavior, receives both olfactory and appetite information, and appears to be able to control movement over the jaw muscles involved in feeding (Roberts, 2012). The middle brain (mesencephalon) is the region particularly involved in the reception and coordination of optic nerve inputs, such as interpretation of motor and visual signals (Baldisserotto et al., 2019; Roberts, 2012). Therefore, we speculate that TiLV infection and neuron damage in this area may be related to behavioral disorders of food and prey seeking, navigating around obstacles in the environment and avoidance of approaching objects. On the other hand, the cerebellum and medulla oblongata are located within the hindbrain (rhombencephalon) and are generally associated with the regulation of locomotion and balance stimuli as well as basal physiology (Baldisserotto et al., 2019; Roberts, 2012). Therefore, it was possible that the strong densities of TiLV in these areas were linked with with erratic swimming or loss of balance. An earlier study also suggested that *Streptococcus agalactiae* infection caused damage in the cerebellum of tilapia and led to the erratic swimming symptoms (Palang, Withyachumnarnkul, Senapin, Sirimanapong, & Vanichviriyakit, 2020). Taken together, localization of TiLV with high density detected in important regions of the brain may contribute to an array of well-described abnormal symptoms, especially neurological disorder signs during the course of infection, and result in high mortality.

Although TiLV was localized in the brain of infected fish (Bacharach et al., 2016; Dong et al., 2017), the mechanism of viral entry to the central nervous system remains unclear. Dong et al (2020) proposed that TiLV causes a systemic infection where the viral particles spread to other organs perhaps via the circulatory system. In this study, the localization of the virus was found in the endothelial cells of the blood vessels, which provided supporting evidence of the virus spread through a hematogenous route resulting in systemic infection (Keller et al., 2003). One of the highlight findings of the TiLV localization in the brain was the gradual increase in the detection of virus-infected cells close to the ventricle. We also noticed that the virus was primarily detected near the periventricular regions, while the area farther away from the ventricle has less detectable infected cells. There has been no report on the infection routes of TiLV into the central nervous system of the fish. However, in case of *Orthomyxovirus*, the possibility that the influenza virus infected the brain of ferrets (Yamada et al., 2012) and chickens (Chaves et al., 2011) through cerebrospinal fluid (CSF) by crossing the blood-CSF barrier has been suggested. Those results have demonstrated that the influenza virus was detected in CSF from infected animals and viral antigen was found in either ependymal cells in the ventricle or choroid plexus epithelial cells (Chaves et al., 2011; Yamada et al., 2012). Similar to the influenza virus invasion, this study revealed that the strong positive signals were dominantly observed in the periventricular regions including the choroid plexus and the ventricular ependymal cells. This suggests that one of the possible routes that the virus can enter to the brain is through the ventricles. Moreover, it is possible that CSF plays an important role in TiLV spreading to broad regions of the brain.

Histopathological alterations in the TiLV infected brain observed in this study included inflammation of the primitive meninges, cell degeneration and blood congestion in multiple regions of the brain, as well as cell aggregation. The latter histopathological feature resembles syncytial cell formation, a pathognomonic change described in liver of TiLV-infected fish (Ferguson et al., 2014), that is occasionally observed in the brain (Behera et al., 2017; Debnath et al., 2020). These damages in the central nervous system might directly or indirectly affect normal functions and homeostasis of the fish brain.

In conclusion, our investigation provided new information on the TiLV distribution in the brain of the experimentally infected Nile tilapia. These findings contribute to the basic knowledge of the disease pathogenesis caused by TiLV and host-pathogen interactions. We also discussed the possible link between TiLV-infected brain regions and abnormal behavioral changes. In addition, we suggested that the ventricles and cerebrospinal fluid are important conduits of TiLV to the brain. However, we have not been able to accurately determine the infected neuronal cells and the spreading pathway. The pathogenesis and distribution of the virus in the central nervous system at the neuronal cellular levels and at various times points of post-infection therefore need to be elucidated in further studies.

## ACKNOWLEDGEMENTS

Nguyen Dinh-Hung acknowledges the scholarship program of Chulalongkorn University for ASEAN and Non-ASEAN countries and the 90^th^ year anniversary of Chulalongkorn University Fund (Ratchadaphiseksomphot Endowment Fund). This study was supported by Mahidol University and TSRI fund (CU FRB640001 01316) from Chulalongkorn University. The authors thank Dr. Vuong Viet Nguyen and Ms. Suwimon Taengphu for their technical assistance.

## CONFLICT OF INTEREST

The authors declare no conflict of interest.

## AUTHOR CONTRIBUTIONS

Conceptualization, H.T.D.; investigation, N.D.H., P.S.; formal analysis, N.D.H., T.K., H.T.D.; methodology, S.S., P.S., N.D.H., H.T.D; supervision; H.T.D., C.R.; writing - original draft, N.D.H., H.T.D.; review & editing, C.R., S.S., T.K., P.K., T.R.R. All authors have read and agreed to the current version of the manuscript.

## Supplementary data

**FIGURE S1.**
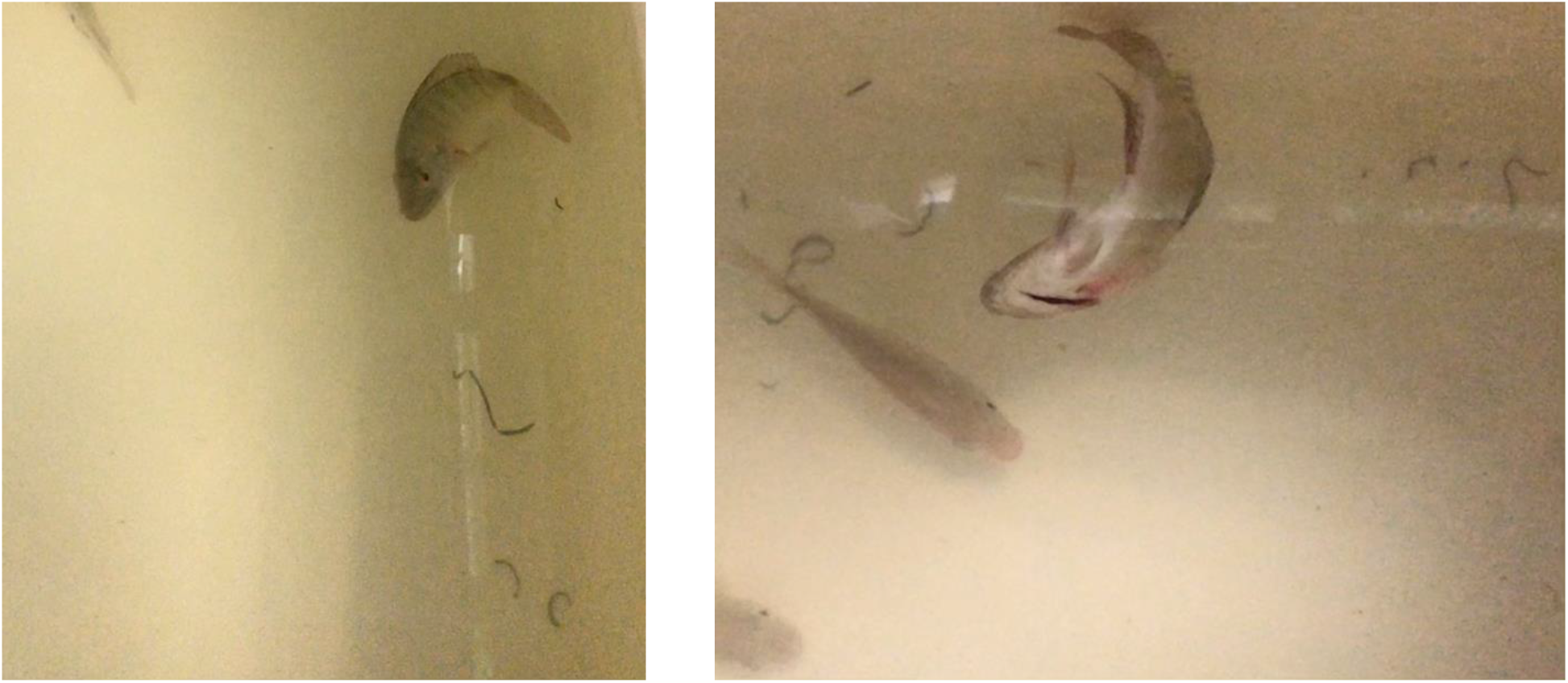
Neurological manifestations were observed in infected fish. The diseased fish displayed erratic swimming, swirling, or loss of balance during 6-9 dpi.

**FIGURE S2.**
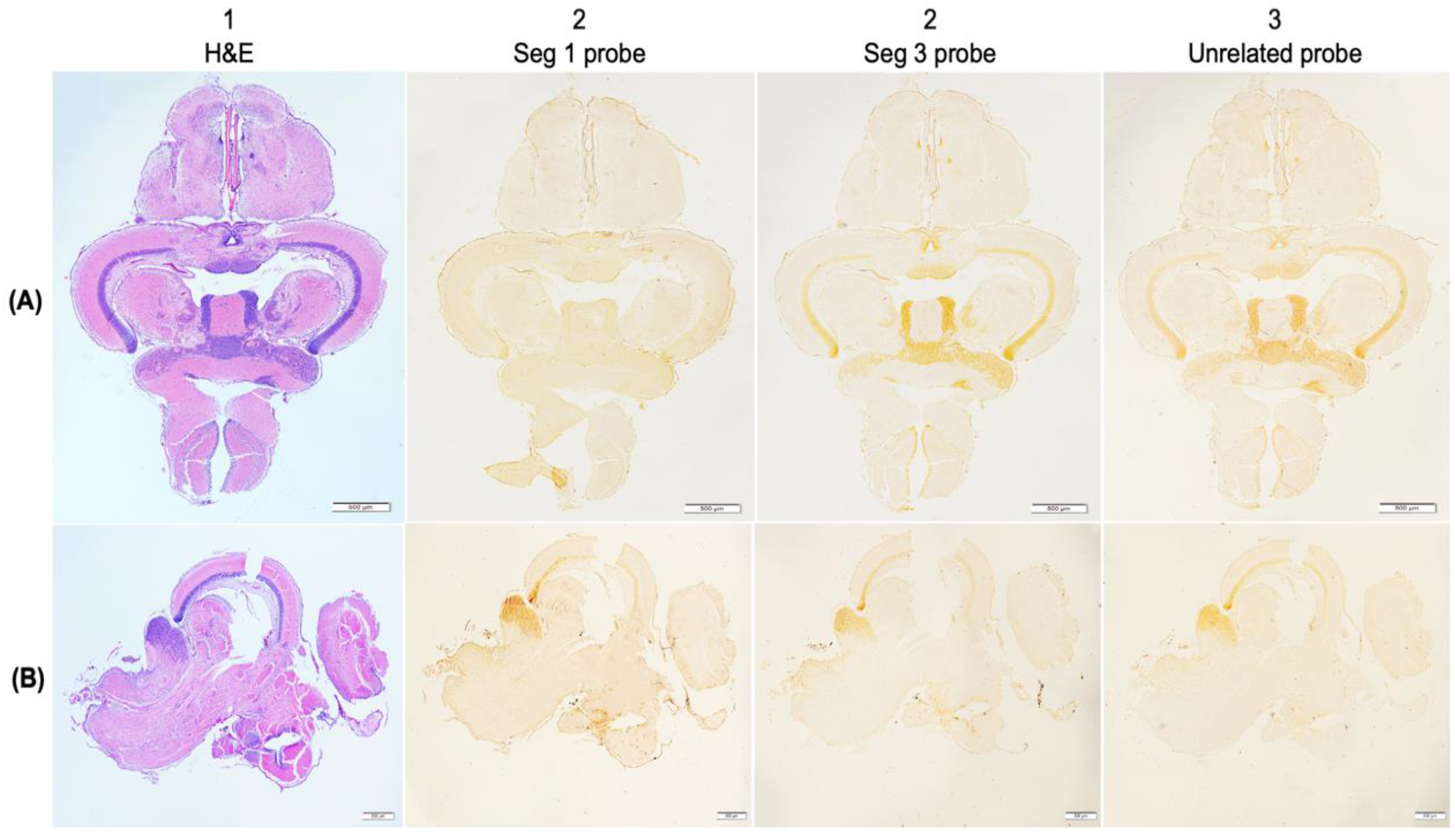
The spatial localization of TiLV in the brain of non-infected fish. (A) and (B): Comparison of horizontal and parasagittal consecutive sections (respectively) were stained with H&E (A.1, B.1), ISH with TiLV probe based on genome segment 1 (A.2, B.2), genome segment 3 (A.3, B.3) and ISH with the unrelated probe as the control (A.4, B.4). No positive signals were found in any brain regions of non-infected fish.

## Notes

### Competing Interest Statement

The authors have declared no competing interest.

